# Historical Areas of Hybridization in Two Striped Emerald Dragonfly Species

**DOI:** 10.64898/2026.10.12.756848

**Authors:** Joseph Caya, Aaron Goodman, Anthony Cognato, Jordy Hernandez, Jessica Ware, Dominic Evangelista

## Abstract

The Hine’s Emerald (Somatochlora hineana) is a federally endangered North American dragonfly, which has experienced significant declines due to anthropogenic forces including human encroachment into its sensitive spring fed wetland habitat and climate change. Previous research indicates hybridization with its ecologically generalist and widespread conspecific, the Clamp-Tipped Emerald (Somatochlora tenebrosa), which may induce hybridization-mediated extinction or genetic swamping. The aim of this paper is to investigate the patterns of hybridization between S. hineana and S. tenebrosa by inferring historical species ranges using Paleoecological niche modeling (PaleoENM), examining genetic structure using haplotype networks, and determining gene flow with introgression tests, shedding light on how, why, when, or where hybridization occurs in these species. Our results reveal evidence of hybridization occurring between S. hineana and S. tenebrosa among populations in Benzie County, Michigan and a newly discovered hybrid individual in Oregon County, Missouri. PaleoENM models show patterns of contact and separation across the Pleistocene, highlighting potential hybridization events throughout periods of glaciation. Finally, Introgression tests reveal evidence for unidirectional hybridization, which may result in genetic swamping of the endangered Hine’s Emerald. Such studies provide essential genetic context for conserving ecologically sensitive and endangered species, identifying conservation priorities, and determining whether hybridization and introgression pose additional threats to the long-term genetic integrity and persistence of endangered populations.

## INTRODUCTION

Anthropogenic-induced climate change drives uneven range shifts in insect species and accelerates the creation of new or altered contact zones that provide the opportunity for interspecific hybridization (Sánchez-Guillén et al., 2016). Shrinking habitats, physiological limits, and species invasions threaten rare species or habitat specialists with limited distributions (Harvey et al., 2023), yet these species face additional pressure from hybridization when confronted by more adaptive and abundant heterospecifics (Armstrong et al., 2025; Gómez et al., 2015; Muhlfeld et al., 2014). Adaptive conspecifics may induce hybridization-mediated extinction or genetic swamping (Chunco, 2014; Sánchez-Guillén et al., 2016; Todesco et al., 2016), thus identifying regions of hybridization and assessing the effects of potential hybridization events are of critical importance (Arce-Valdés & Sánchez-Guillén, 2022). Hybrid zones are often formed at parapatric boundaries between taxa or within the distribution of co-occurring sympatric taxa (Kawakami & Butlin, 2012), yet the ability to verify the presence of hybrid zones has historically been limited to morphological analyses of field samples. Today, species distribution modeling – powered by occurrence and environmental datasets – is a powerful tool that when fully utilized can identify species’ overlapping clines (Taylor et al., 2015). Genetic techniques supplement geographic observations with strategies such as phylogenetic reconstruction (Kawakami & Butlin, 2012), introgression (ABBA-BABA tests) (Martin et al., 2015), and haplotype networking (Maresova et al., 2019) that may generate insights about the origins, movement, and effects of hybrid zones on species. Hybridization in the context of conservation has been explored thoroughly for reptiles, birds, and mammals (Bonnald et al., 2021; Doniol-Valcroze et al., 2024; Morales-Rozo et al., 2017; Shurtliff, 2013) as well as plant species (Cozzolino et al., 2006; Muniz et al., 2022), however there are limited studies exploring endangered or threatened arthropods, including dragonflies.

Dragonfly (Odonata) biodiversity embodies incredible dispersal capability, habitat specificity, reproductive strategy, and sensitivity to environmental gradients which can facilitate study of hybrid zones unlike other animal groups (Bybee et al., 2016; Corbet, 1999; Córdoba-Aguilar, 2008). Dragonflies present an ideal system to understand the role of hybridization within conservation, with an evolutionary history extending back ∼325 mya and ∼6400 extant species currently described, whose distribution range from single-stream endemics to transoceanic migrants with life cycles connecting freshwater and terrestrial habitats (Dijkstra, 2025; Newton et al., 2026). As indicators of freshwater ecosystem health and sentinels of climate change, odonates are among the most valuable insect groups for evaluating conservation success (Grant & Samways, 2007; Samways et al., 2025), thus investigations into hybridization are well within the scope of current conservation efforts and may lead to impactful discoveries for odonate conservation. At present, studies of hybridization in odonates have been focused on damselflies (Zygoptera) (Hayashi et al., 2005; Sánchez-Guillén et al., 2012; SÁNCHEZ-GUILLÉN et al., 2005; Solano et al., 2018; Tynkkynen et al., 2008; Wellenreuther et al., 2018), with minimal work on dragonflies (Anisoptera) (Solano et al., 2018).

The dragonfly genus *Somatochlora* stands out due to anecdotal evidence suggesting several instances of heterospecific hybridization (Bick & Bick, 1981; E. M. Walker, 1925), including *S. albicincta* and *S. hudsonica* individuals hybridizing with *S. sahlbergi* in the Northern Yukon (S. G. Cannings & Cannings, 1997), as evidenced by intermediate morphologies.

Although these three species are observed to have specialized habitat preferences (S. Cannings & Cannings, 1985; Catling, 2003), hybridization seems to occur whenever they come into contact, causing difficulty in species-level identification. Among *Somatochlora* lies the only federally endangered dragonfly species, Hine’s Emerald (*Somatochlora hineana*), whose status is attributed to the loss of its rare habitat and subsequent decline over the past century (USFWS, 2001). Historical records indicate populations of Hine’s Emerald once persisted across a wider range, including Alabama, Ohio, and Indiana, yet their specialized habitat no longer exists in these states, leaving a handful of isolated populations in Missouri, Illinois, Wisconsin, and Michigan (Walker et al., 2020). Populations of Hine’s Emerald require fens fed by calcareous seepage flow (Cashatt & Vogt, 2001; Vogt & Cashatt, 1994), the presence of crayfish burrows for sheltering nymphs (Pintor & Soluk, 2006), and nearby meadows for adult foraging (Foster & Soluk, 2006). Habitat loss remains a primary concern for agencies protecting Hine’s Emerald (USFWS, 2024), yet the discovery of new populations in regions lacking one or several of the aforementioned characteristics suggests Hine’s Emerald may be less restricted than previously thought (Craves et al., 2022; Kessler et al., 2026).

Recent work by Hernandez and Cognato (2024) provides the first molecular evidence of heterospecific mating and introgression within *S. hineana* + *S. tenebrosa*, *S. kennedyi* + *S. forcipata* and *S. calverti* + *S. provocans* + *S. filosa*. Phylogenies generated using mitochondrial DNA recover unresolved groupings among these species. However, phylogenies generated using nuclear DNA resolve species-level relationships with high support, suggesting high degrees of recent hybridization. Hernandez and Cognato (2024) hypothesize that increased sexual pressure to mate with heterospecifics result in the dilution of genotypes and that prezygotic barriers (i.e., male cerci) are much less pertinent to reproductive isolation than previously thought. It is unknown if this asymmetrical mitochondrial gene flow, as observed in other insects (e.g. Cognato et al., 1999) is prevalent in all areas of *S. hineana* and *S. tenebrosa* sympatry (Hernandez & Cognato, 2024). The lack of information or documented records of hybridization in such a fragile species demands attention from researchers, which we aim to resolve through our study.

In order to address the critical knowledge gaps regarding hybridization among Hine’s Emerald populations, we employ paleoecological niche modeling (paleoENM) and genomic-based introgression and phylogenetic tests to shed light on its history of interaction with the Clamp-Tipped Emerald (*Somatochlora tenebrosa*). We produce paleoniche distributions of both species from the Last Interglacial period (ca. 130 Ka) to the present, elucidating historical patterns of sympatry and potential hybridization. We supplement our geographic predictions with a reconstructed anchored-hybrid enrichment (AHE) phylogeny and haplotype networks based on the mitochondrial (COI) gene for both species. Additional introgression tests based using single nucleotide polymorphism (SNP) data test for admixture between species.

Based on Hernandez and Cognato’s (2024) results of introgression between Hine’s Emerald and the Clamp-Tipped Emerald, we expect to replicate evidence of hybridization in our haplotype networks and introgression tests. Signals of hybridization should be reflected in the paleo niche models by overlapping ranges, which may provide insights on the history of hybridization among these dragonflies. Moreover, range models and genetic tests will help to evaluate the patterns in hybridization among these species, allowing us to shift our focus to the potential outcomes in persistence or resilience of the Hine’s Emerald as a result of hybridization.

## METHODS

### Taxon Sampling

We obtained aliquots of DNA from the same specimens of *S. hineana* and *S. tenebrosa* previously used in Hernandez and Cognato (2024) which were field-collected and/or sourced from museum collections. In total, 21 specimens of *S. hineana* and 20 of *S. tenebrosa* were included.

### DNA Extraction and Sequencing

DNA was extracted from a meso- or meta-leg with the DNeasy Blood and Tissue Kit (Qiagen, Hilden, Germany) following manufacturer protocols. We quantified DNA yield using a Qubit 4 fluorometer, and sent our samples to RAPID Genomics (Gainesville, Florida) for library preparation and sequencing. We amplified reads using modified Anchored Hybrid Enrichment (AHE) probes first implemented in Bybee et al. (2021). We acquired reads using a 92 loci (20kb) probe set (Goodman et al., 2023). We acquired additional reads consisting of the full 1,306 probe set (500kb) (Bybee et al., 2021) from outgroups generated from (Goodman et al., 2025). Sampled outgroups included select genera from ingroup Libelluloidea (Corduliidae, Libellulidae, Synthemistidae, Macromiidae), and outgroup Cavilabiata (Cordulegastridae, Neopetaliidae, Chlorogomphidae).

### Phylogenetic Analysis

We assembled AHE reads, assigned orthology, and generated alignments following Goodman et al. (2023) and Goodman et al. (2026), and trimmed loci from our 500kb specimens to match the number of loci retained across *Somatochlora* samples. We generated multiple sequence alignments for each locus using the ‘MAFFT-linsi’ algorithm in *MAFFT v.7.475* (Katoh & Standley, 2013), and trimmed alignments using a 0.75 threshold cutoff using *trimAI v1.2* (Capella-Gutiérrez et al., 2009). We concatenated the alignment using *FASconCAT v1.11* (Kück & Meusemann, 2010), and generated an initial partitioning scheme using relaxed clustering with the model fixed to GTR + G for each subset in *IQtree v2.1.*3 (Minh et al., 2020). We then selected the best nucleotide substitution model for each subset in the partitioning scheme using ModelFinder and estimated a maximum likelihood tree (ML). We estimated branch support using 1,000 ultrafast bootstrap replicates (UFboot) in *IQtree v2.1.3* (Guindon et al., 2010; Kalyaanamoorthy et al., 2017). Finally, to assess the degree of incomplete lineage sorting, we performed a coalescent-based species tree estimation in *ASTRALIII v5.6.1* (Zhang et al., 2018) and estimated branch support using local posterior probabilities (LPP) (Mirarab & Warnow, 2015). We identify nodes of high support possessing ML and SH-aLRT values >95 and LPP values >0.95. We rooted the tree using our outgroups.

### Haplotype Analysis

To assess geographic variation among individuals of *S. hineana* and *S. tenebrosa*, we reconstructed minimum spanning networks using PopArt version 1.7 (Leigh et al., 2015) with the minimum spanning networks algorithm (Bandelt et al., 1999). We explored genetic variation among species using the Cytochrome Oxygenase I (COI) gene, which was among the loci amplified in our AHE dataset. We performed three separate analyses, in which we reconstructed haplotype networks for *S. tenebrosa*, *S. hineana*, and a combined network of both species. Due to gaps in the sequences, we removed four specimens (SOM 111, SOM 139, SOM 147, SOM55) prior to analysis.

### Introgression Tests

Using only our *Somatochlora* samples, we mapped raw AHE reads from all individuals to a soft-masked genome assembly of *Somatochlora meridionalis* (Genoscope NCBI) using *bwa* (Li & Durbin, 2009) and sorted alignments with *samtools* (Danecek et al., 2021). We called variants using *bcftools* (Danecek et al., 2021). Previous research has suggested hybridization occurring between *S. hineana* and *S. tenebrosa* (Hernandez & Cognato, 2024), specifically within two samples collected within Benzie County Michigan (SOM 11 & 12 in Cognato and Hernandez 2025), as well as a third sample (SOM 159), which we recovered as a hybrid in preliminary haplotype analyses. We tested for evidence of introgression and gene flow between our three putative hybrid individuals and *S. hineana, S. tenebrosa* using an ABBA-BABA test from *Dsuite* v0.5 r57 (Malinsky et al., 2021), which is optimized for more evolutionarily recent lineages.

We performed pairwise configurations of the ABBA-BABA test by swapping the relatedness of *S. hineana* and *S. tenebrosa* in relation to the putative hybrid. Outgroups consisted of 51 individuals of *Synthemis eustalacta* (See Goodman et al., 2026).

### Occurrence Records

We acquired occurrence records of adult *S. hineana* and *S. tenebrosa* from the Global Biodiversity Information Facility using the R package ‘spocc’ (Owens et al., 2025) along with our records from Hernandez and Cognato (2024). We selected occurrences from GBIF possessing preserved museum samples and research grade observations, which include verified latitude and longitude coordinates, a photograph of the sighting, observation date, and at least ⅔ agreement on species identification by the community. While our records include other extirpated populations of *S. hineana*, we excluded a single occurrence from Jackson County, Alabama due to the fact this individual was never associated with an active breeding population (Vogt & Cashatt, 1994). We then filtered occurrences by removing records with duplicate coordinates, and using the R package ‘spThin’ (Aiello-Lammens et al., 2015), we spatially thinned occurrences within 5 km to account for sampling bias.

### Environmental Data

All analyses were conducted using the statistical programming language R v. 4..6.0 (R Core Team, 2021), and all data are in a latitude/longitude coordinate system using WGS84 datum. We acquired averaged sets of environmental predictor variables consisting of strictly climatic data. We acquired environmental rasters at 2.5 arc-second resolution (∼ 5 km at the equator) for both modern settings. We downloaded CHELSA’s ‘Anthropocene’ bioclimatic dataset (1979–2013) consisting of 19 bioclimatic variables extrapolated from monthly minimum and maximum temperature, humidity, mean, and coefficient of variation of annual solar radiation (Karger et al., 2023) (herein referred to as our ‘modern day’ dataset). We downloaded PaleoClim’s bioclimatic datasets ranging from the Last Glacial Maximum (120 ka) to the late Holocene (0.3 Ka) (Brown et al., 2018). Variables bio08, bio09, bio18, and bio19 were removed from all datasets after visual inspection of interpolation discontinuities in our study extent (Booth, 2022).

### Ecological Niche Modeling

Prior to modeling, we defined a 300km buffer around occurrences to represent the dispersal limitations of *Somatochlora* species, which was used to sample background points. We performed additional thinning of occurrences to the resolution of our environmental variables to reduce the effects of sampling bias and artificial clustering (Veloz, 2009) using the ENMtools package (Warren et al., 2010) and extracted the environmental values from remaining points.

We used these values to calculate correlations between variables using the ‘vifstep’ function in the ‘usdm’ package (Naimi & Naimi, 2017) and filtered out variables with a VIF threshold of 7. Variable selection with ‘vifstep’, retained bio3 or bio4 depending on the sequence of variables eliminated. Bio4 was retained preferentially because it is derived from a single climatic variable, thus we expect the interpretation of bio4 to be more reliable across paleoclimate predictions.

To model the distribution of *S. hineana* and *S. tenebrosa*, we used the presence-background algorithm Maxent v3.4.4 (Phillips et al., 2017), one of the top-performing models for fitting ENMs with background data (Valavi et al., 2022). We used the R Package ‘ENMeval’ 2.0.5 (Kass et al., 2021) for model building, parameterization, evaluation with different complexity settings, and reporting of results. Due to *S. hineana’s* disjunct distribution, checkerboard partitioning was used to maintain environmental similarity between the training and validation datasets. We clamped our models by omitting ranges of environmental data which fall outside of our training data to prevent projection of our model to non-analogue climates. We tuned model complexity to find optimal settings (Radosavljevic & Anderson, 2014; Warren & Seifert, 2011). For tuning, in order of increasing complexity, we chose the feature classes linear (L), linear quadratic (LQ), hinge (H), and linear quadratic hinge (LQH), as well as regularization multipliers one through four (higher numbers penalize complexity more).

We assessed our model using averages of threshold-dependent (omission rate) and threshold-independent (AUC) discrimination performance metrics calculated on withheld validation data, (Warren & Seifert, 2011). We assessed the 10-percentile omission rate, which sets a threshold as the lowest suitability value for occurrences after removing the lowest 10% suitability values (Kass et al., 2021; Radosavljevic & Anderson, 2014). Validation AUC is a measure of discrimination accuracy that can be used to make relative comparisons between ENMs with different settings fit on the same data (Lobo et al., 2008; Radosavljevic & Anderson, 2014). Optimal models for both *Somatochlora* species were selected by AICc deltas of 0.

Our ecological niche models are based on present-day occurrence records of *S. hineana* and *S. tenebrosa* and environmental predictor variables from CHELSA’s ‘Anthropocene’ dataset. Habitat suitability was then mapped in the geologic past by using variables from the PaleoClim dataset (Brown et al., 2018). We generated habitat suitability predictions within the following geologic periods: late-Holocene (Meghalayan: 0.3–4.2 ka), mid-Holocene (Northgrippian: 4.2–8.326 ka), early Holocene (Greenlandian: 8.326–11.7 ka), Younger Dryas Stadial (12.9-11.7ka), Bølling-Allerød (14.7-12.9 ka), Heinrich Stadial (17.0-14.7 ka), Last Glacial Maximum (21 ka) and Last Interglacial (ca. 130 ka). Predictions were estimated using a new study extent, generated as a bounding box spanning the eastern United States delimited by the following unprojected coordinates: −100° to 57° longitude, 28° to 51° latitude. We chose this new study extent for our predictions to estimate potential areas of sympatry between *S. hineana* and *S. tenebrosa*. Maxent raw predictions were transformed to a scale of 0–1 to approximate probability of occurrence using the ‘cloglog’ transformation (now referred to as our continuous prediction) (Phillips et al., 2017). We also generated a threshold prediction to map areas of congruency in PaleoClim layers, calculated from the 10-percentile omission rate from our model evaluation. All habitat suitability predictions were exported as rasters for visualization in ArcGIS Pro.

### Niche Overlap

In order to assess the niche overlap among *S. hineana* and *S. tenebrosa* in each time period, we normalized our continuous habitat suitability predictions to calculate pairwise Schoener’s D (Schoener, 1968) and Hellinger’s I (Warren et al., 2008) indices, where both indices range from 0 (no overlap) to 1 (complete overlap). Schoener’s D assesses the absolute differences in the relative distribution of suitability across geographic space, whereas Hellinger’s I assesses the shared component of the two suitability distributions. Both metrics accomplish the same goal with different mathematical approaches, providing complementary measures of niche overlap.

## Results

### Phylogenetic Analysis

We recover the validity of *Somatochlora hineana* and *S. tenebrosa* with high support (BS: 100, PP: 99), though intra-specific relationships within these species were poorly supported (BS: 26–83, PP: 0.39–0.77). We observed minimal phylogeographic structuring among populations of either species **(Supplemental Fig. S1)**. Furthermore, we observed high degrees of topological discordance between our ASTRAL and ML phylogenies within species **(Supplemental Fig. S2)**.

### Haplotype Analysis

We observed minimal geographic structuring when specimens were coded by haplotype and mapped. Both *S. hineana* and *S. tenebrosa* networks, coded by haplotype, recovered one dominant haplotype each (H2 and H3 respectively) present across most sampled regions, with derived variants dispersed throughout the species’ ranges **(Fig. 1a**, **1b)**. In the combined network, coded by species, the haplotype network showed a *S. tenebrosa* haplotype within the dominant *S. hineana* cluster, as well as variants diverging from the *S. hineana* network **(Fig. 1e)**.

**Figure 1.**
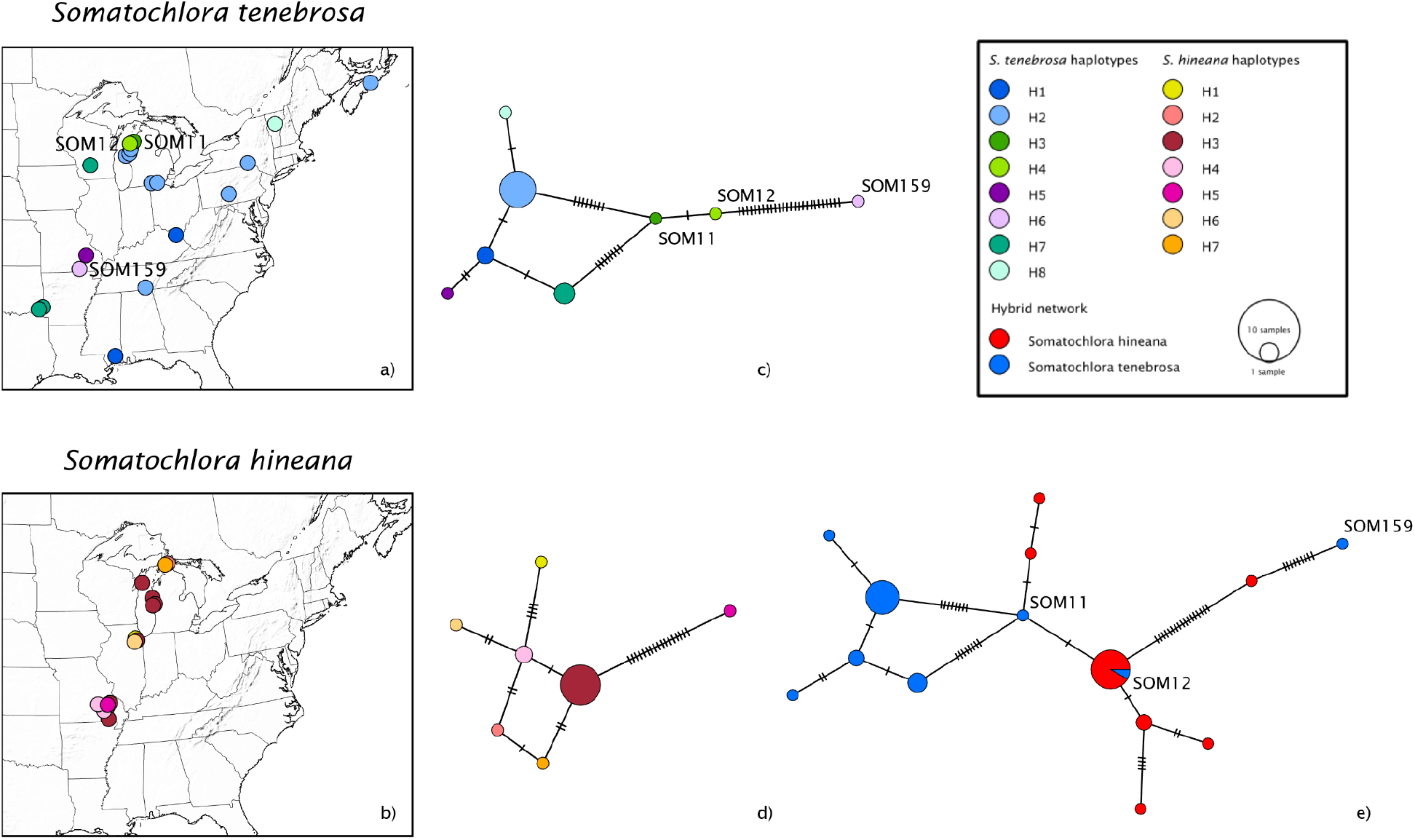
Maps of DNA samples for *S. tenebrosa* (a) and *S. hineana* (b). Points represent one record each and are displaced up to 50km for visualization. Also shown are haplotype networks for *S. tenebrosa* (c)*, S hineana* (d), and a combined network (e). Putative hybrids SOM11 and SOM12 from Hernandez and Cognato (2024) in addition to our new hybrid SOM159 are annotated.

### ABBA-BABA Test

All three configurations of *Somatochlora* species and putative hybrids returned significant D-statistics (D = 0.137–0.302, P ≤ 0.00142, Z = 3.19–8.84; **Table 1**). Our three putative hybrid individuals shared derived alleles most frequently with *S. tenebrosa* (BBAA = 374.82), indicating a closer relationship to *S. tenebrosa* than *S. hineana*. As sister taxa are expected to share the most derived alleles, we analyzed introgression under this arrangement. *S. hineana* shared derived alleles with the hybrids (ABBA = 264.76) more frequently than with *S. tenebrosa* (BABA = 201.01), yielding signatures of introgression (D = 0.137, Z = 3.19, P = 0.00142, f4 = 0.139; **Table 1**).

**Table 1.** ABBA–BABA test results generated with Dsuite Dtrios. We compared putative hybrid individuals with *S. hineana* and *S. tenebrosa* with *Synthemis eustalacta* as the outgroup. Each row represents a different tree configuration under the four-taxon topology ((P1, P2, P3), Outgroup). D-statistic represents minimum Patterson D values across the three configurations. Z-score assess significance of deviation from zero. f4-ratio estimates the proportion of admixture. BBAA, ABBA, and BABA are counts of derived allele site patterns.

| Tree | P1 | P2 | P3 | Dstatistic | Z-score | p-value | f4-ratio | BBAA | ABBA | BABA |
| --- | --- | --- | --- | --- | --- | --- | --- | --- | --- | --- |
| 1 | hineana | hybrid | tenebrosa | 0.301856 | 8.84222 | 2.30E-16 | 0.511151 | 264.763 | 374.824 | 201.006 |
| 2 | tenebrosa | hybrid | hineana | 0.136884 | 3.19078 | 1.42E-03 | 0.139097 | 374.824 | 264.763 | 201.006 |
| 3 | hineana | tenebrosa | hybrid | 0.172082 | 5.31016 | 1.10E-07 | 0.233081 | 201.006 | 374.824 | 264.763 |

### Ecological Niche Modeling

Optimal model settings for *S. hineana* and *S. tenebrosa* exhibited robust results based on validation AUC, yet 10-percentile omission differed between species (0.24 and 0.09 respectively). Based on the results of the collinearity analysis, we used the following five predictor variables to build both models: Mean temperature of the warmest quarter (bio10), precipitation of the wettest month (bio13), precipitation seasonality (bio15), mean diurnal range (bio02), and temperature seasonality (bio04).

The optimal model for *S. hineana* was complex, with linear, quadratic, and hinge feature classes and a regularization multiplier of 1 (low complexity penalty). Our mean validation AUC was high (0.81), and a 10% omission rate that was high (0.24) relative to its expected value of 0.1. Furthermore, the optimal model possessed an AICc of 1483.67, a delta AICc of 0, and 22 non-zero lambda values, indicating high overfitting of model prediction (**Table 2**). Precipitation of the wettest month (bio13) possessed the highest variable importance (86%), followed by temperature seasonality (bio04) (5.8%), while mean temperature of the warmest quarter (bio10), and precipitation seasonality (bio15), and mean diurnal range (bio02) possessed < 4% (**Supplemental Fig. S3**). Mean temperature of the warmest quarter (bio10), precipitation seasonality (bio15), mean diurnal range (bio02), and temperature seasonality (bio04) expressed quadratic relationships with suitability, containing jagged optimal ranges most likely due to the disjunct distribution of *S. hineana* populations. Precipitation of the wettest month (bio13) expressed a hinged, linear relationship with suitability. The greatest suitability occurs at 19°C and 25°C for mean temperature of the warmest quarter (bio10), <90mm for precipitation of the wettest month (bio13), 12% and 30% for precipitation seasonality (bio15), 5-9°C for mean diurnal range (bio02), and 8.5-9.5°C for temperature seasonality (bio04) (**Supplemental Fig. S3**).

**Table 2.** Sample size of occurrence localities before and after spatial thinning and maxent ENM settings for *Somatochlora hineana* and *Somatochlora tenebrosa*. Settings shown are feature class (features). Statistics shown are mean validation AUC (AUCval), mean omission rates for 10 percentile training values (OR10), Akaike Information Criterion (AICc), and delta Akaike Information Criterion (delta.AICc), and number of nonzero coefficients (ncoef).

| Species | Raw occurrences | Thinned occurrences | Features | Regularization | AUC<sub>val</sub> | AICc | delta.AICc | OR10 | ncoef |
| --- | --- | --- | --- | --- | --- | --- | --- | --- | --- |
| *Somatochlora hineana* | 147 | 68 | LQH | 1 | 0.81 | 1483.67 | 0 | 0.24 | 22 |
| *Somatochlora tenebrosa* | 1507 | 859 | LQH | 1 | 0.83 | 20206.26 | 0 | 0.09 | 28 |

The optimal model for *S. tenebrosa* was complex, with linear, quadratic, and hinge feature classes and a regularization multiplier of 1 (low complexity penalty). Our mean validation AUC was high (0.83), and a 10% omission rate that was low (0.098) relative to its expected value of 0.1. Furthermore, the optimal model possessed an AICc of 20206.26, a delta AICc of 0, and 28 non-zero lambda values, indicating high overfitting of model prediction (**Table 2**).

Precipitation seasonality (bio15) possessed the highest variable importance (72%), followed by mean temperature of the warmest quarter (bio10) (16%) and temperature seasonality (bio04) (8.5%), while precipitation of the wettest month (bio13) and mean diurnal range (bio02) possessed < 3% (**Supplemental Fig. S4**). Mean temperature of the warmest quarter (bio10), mean diurnal range (bio02), and temperature seasonality (bio04) expressed quadratic relationships with suitability. Precipitation of the wettest month (bio13) and precipitation seasonality (bio15) expressed linear relationships with suitability. The greatest suitability occurs at 18-26°C for mean temperature of the warmest quarter (bio10), >75mm for precipitation of the wettest month (bio13), <25% for precipitation seasonality (bio15), 8°C for mean diurnal range (bio02), and 8°C for temperature seasonality (bio04) (**Supplemental Fig. S4**).

Continuous predictions over the study extent for our ‘Anthropocene’ environmental variables revealed areas of high suitability in the Great Lakes, Ozarks, and Northeast regions of the United States for *S. hineana* (**Fig. 2a**). Predictions of *S. tenebrosa* revealed areas of high suitability in the eastern Great Lakes, Ozarks, and East Coast regions of the United States (**Fig. 2b**). Areas of congruency mapped with our thresholded predictions emerge anywhere the *S. hineana* model fitted above the threshold, with the exception of our ‘Anthropocene’ variables (**Fig. 3**). Its predicted range is contained within miniscule regions along the east coast until the late holocene, where greater areas of high suitability emerged across the Northeast, Great Lakes, and Ozarks regions.

**Figure 2.**
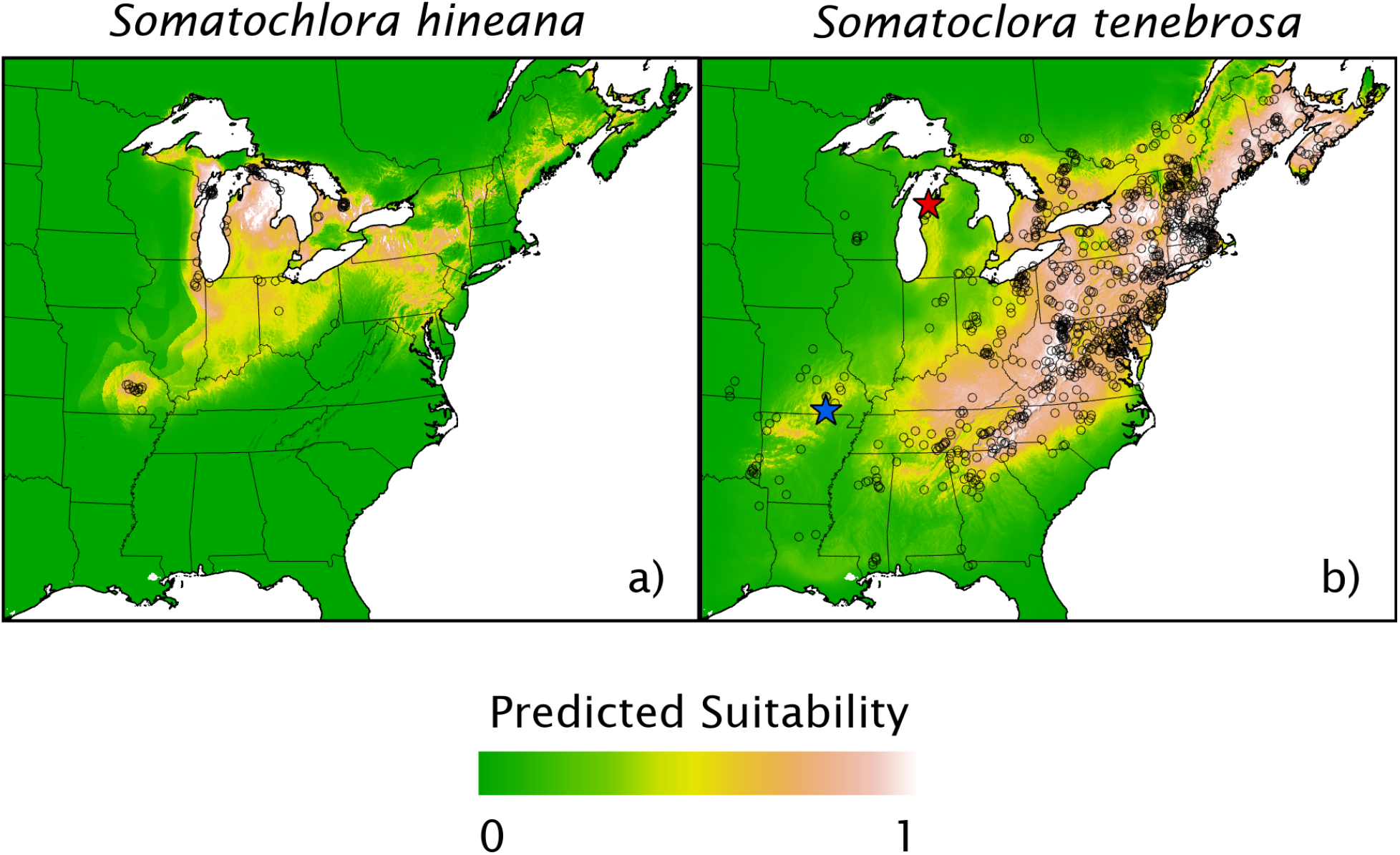
Continuous maxent predictions for *S. hineana* and *S. tenebrosa* projected using our ‘Anthropocene variables’ and range of thinned occurrences for *S. hineana* and *S. tenebrosa* throughout North America. Circles represent one occurrence record, a red star represents our two putative hybrids in Benzie County, Michigan (SOM11 and SOM12), and a blue star represents our new hybrid individual in Oregon County, Missouri (SOM159). Predictions were derived from the optimal model using the criterion of AUCtest values being the highest, and 10% omission rate being the lowest. Predictions were transformed using the ‘cloglog’ function in Maxent v3.4.4 (Phillips et al., 2017), in which raw values are converted to a range of 0–1 to approximate a probability of occurrence. Brighter colors (white, orange, yellow) indicate areas of higher suitability (higher probability of occurrence), while green indicates areas of lower suitability (lower probability of occurrence)

**Figure 3.**
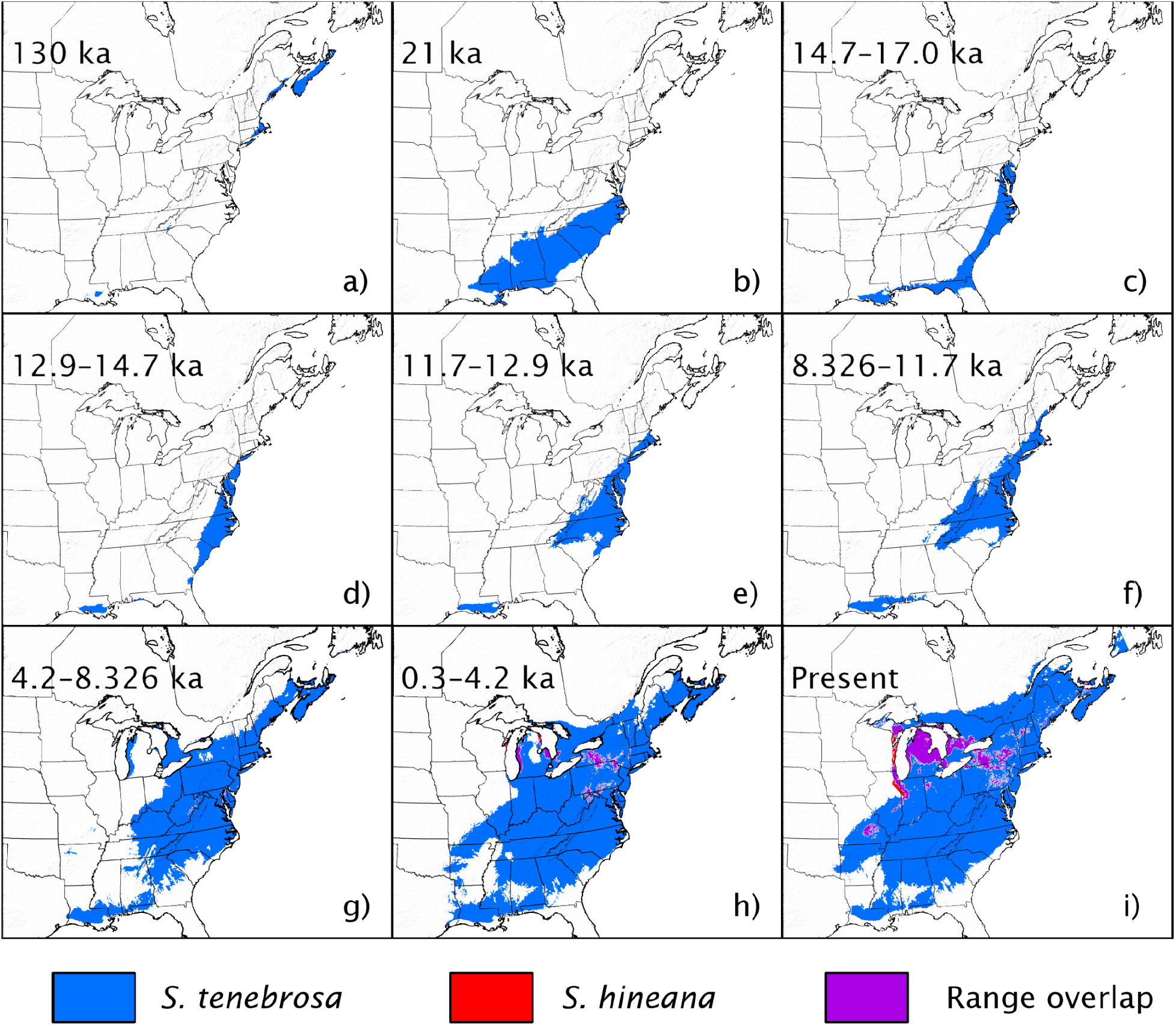
Binary maxent predictions for *S. hineana* and *S. tenebrosa* projected onto paleoclimatic layers in the following order: The Last Interglacial (LIG) (130 ka) (a), Last Glacial Maximum (LGM) (21 ka) (b), Heinrich Stadial (14.7–17.0 ka) (c), Bølling-Allerød (12.9–14.7 ka) (d), Younger Dryas Stadial (11.7–12.9 ka) (e), early Holocene (Greenlandian: 8.326–11.7 ka) (f), mid-Holocene (Northgrippian: 4.2–8.326 ka) (g), late-Holocene (Meghalayan: 0.3–4.2 ka) (h), and Anthropocene (i). Predictions were derived from the optimal model using the criterion of AUCtest values being the highest, and 10% omission rate being the lowest. Predictions were calculated from the 10-percentile omission rate from our model evaluation, which visualizes our most conservative estimation of presence-absence.

### Niche Overlap

Our tests for niche overlap based on continuous suitability predictions showed distinct periods of overlap throughout the Quaternary (**Fig 4**). The Schoener’s D (**Fig 4a**) tests showed high degrees of overlap between *S. hineana* and *S. tenebrosa* in the Heinrich Stadial and Bolling-Allerod (0.36-0.39), with greater values in the Mid Holocene, Late Holocene, and present (0.40-0.47). Hellinger’s I (**Fig 4b**) yielded similar results, showing high degrees of overlap during the Last Interglacial, Heinrich Stadial, and Bolling-Allerod (0.58-0.62), with greater values in the Mid Holocene, Late Holocene, and present (0.66-0.74).

**Figure 4.**
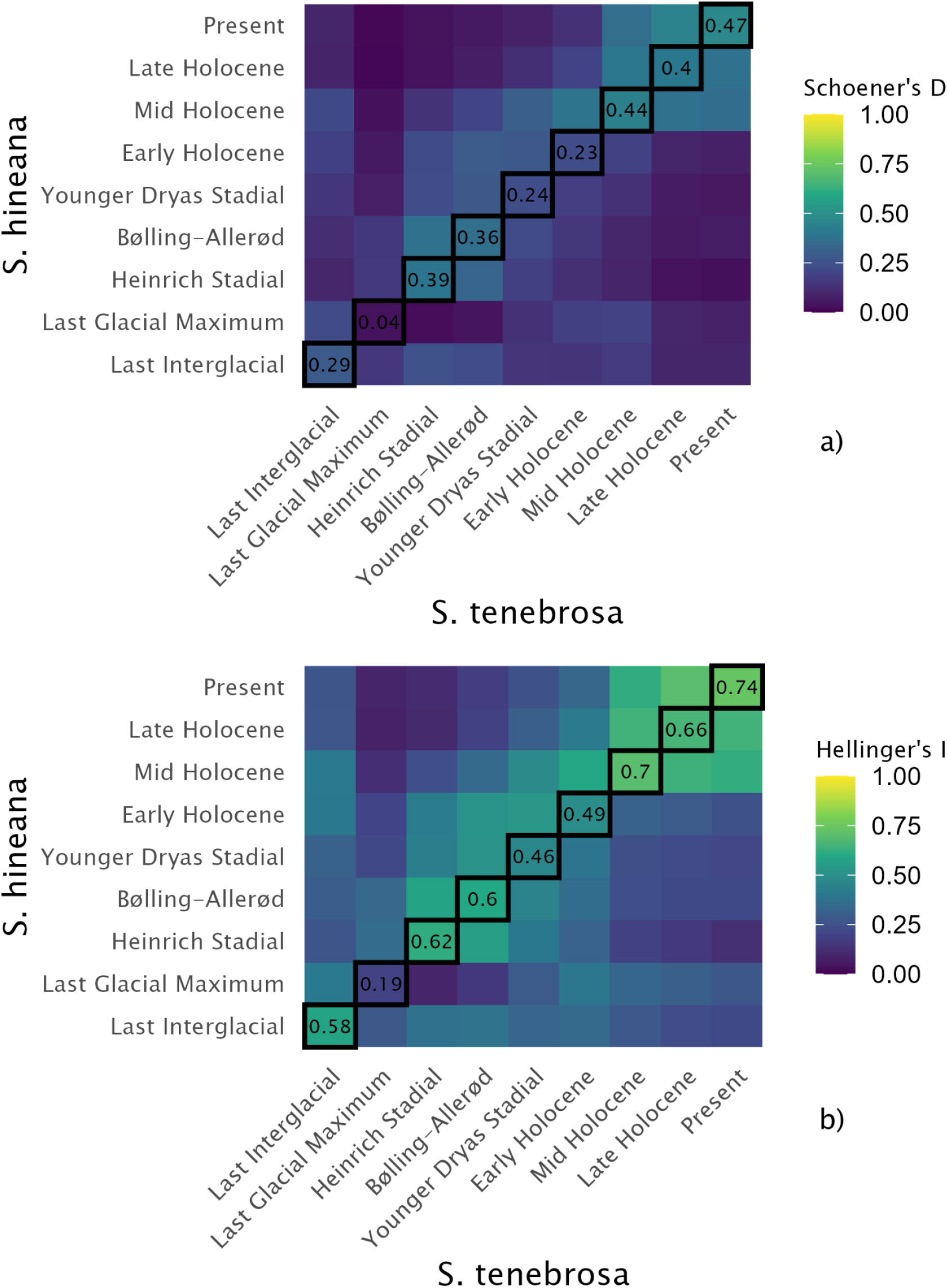
Schoener’s D and Hellinger’s I indices measure the overlap of continuous Maxent predictions for *S. hineana* and *S. tenebrosa*. Schoener’s D assesses the absolute differences in the relative distribution of suitability across geographic space, whereas Hellinger’s I assesses the shared component of the two suitability distributions.

## Discussion

### Ecological Niche Models

Both models shared similar responses to mean temperature of the warmest quarter (bio10), mean diurnal range (bio02), and temperature seasonality (bio04), yet *S. hineana* expresses steep cutoffs in suitability for bio04. These results suggest *S. hineana* is climatically sensitive, yet it is unclear if this is a byproduct of the species being tied to calcareous fens (Kessler et al., 2026). On the other hand, the abundance of occurrence records and large suitability range support anecdotal claims of *S. tenebrosa* being a common, generalist species (Dunkle, 2000; White et al., 2014). Both models expressed precipitation (bio13 and bio15) as a highly important variable, however we observe contrasting patterns among both species. Within *S. tenebrosa*, we observed increases in suitability with precipitation of the wettest month (bio13) and decreases with precipitation seasonality (bio15). We hypothesize that increased rainfall during its peak flight period (May-Oct) opens more niche space due to the increased availability of streams. Furthermore, we hypothesize that the species prefers regions where precipitation-fed streams are non-ephemeral throughout its flight period. Within *S. hineana*, we observed decreases in suitability with precipitation of the wettest month (bio13) and peaks with precipitation seasonality (bio15). We hypothesize that increases in rainfall during its peak flight period (May-Sept) results in decreased oviposition habitat due to the inundation of spring-fed calcareous fens that alters water chemical properties and may affect flora adapted to shallows (Almendinger & Leete, 1998; Almendinger* & Leete, 1998). Calcerous fens are characterized as seasonally dry, which may explain the quadratic relationship with suitability in precipitation seasonality (bio15) (Dunkle, 2000). Finally, several of *S.* hineana’s response curves pertaining to temperature (bio10, bio2) were observed to be bimodal, possibly due to the disjunct distribution of occurrences; an observation occurring among other disjointly distributed odonate taxa (Goodman et al., 2024).

### Modern-Day Distribution

Our continuous prediction of *S. hineana* over modern day variables revealed patterns generally consistent with Kessler et al. (2026), who generated high-resolution suitability maps for the species. Our projections include additional areas of suitability in the Northeast, although there are no recorded observations of *S. hineana* in this region, meaning the model may represent a relictual distribution or extrapolation beyond the dispersal capability of the species. Suitability for *S. tenebrosa* over modern day variables is widespread, with areas of high suitability emerging throughout the Appalachian range and greater Eastern U.S. There are yet many occurrence records that fall outside regions of predicted suitability, which is expected given the supposed generalist habit of *S. tenebrosa*. It should also be noted that in Kessler et al.’s (2026) modeled distributions of *S. hineana*, landscape characteristics (WetSoils, NLCD) emerged as far more reliable predictors of habitat suitability than climatic variables, yielding higher model performance than with climatic variables alone. Kessler et al. (2026) additionally obtained higher model performance with regional models than nationwide models, highlighting the challenges associated with modeling this sparsely distributed species. Given these insights, information gained through our paleodistributions should be interpreted purely through the lens of large-scale climate trends.

### Paleo Distribution

Throughout our Paleoclim time slices, both our continuous and thresholded predictions show the species moving north-south in accordance with glacial advancement and retreat with clear patterns of contact and separation. Previous research has observed genetic signatures of northward dispersal into post-glacial landscapes within *S. hineana* (Purdue et al., 1999; J. Walker et al., 2020). Previous niche models of odonates have suggested their use of refugia during periods of glaciation (Abbott et al., 2022), however this hypothesis has not been explicitly tested in odonates. Niche overlap tests showed distinct periods of intersection and separation, indicating the possibility these species have been in contact throughout the past. Since we observe high overlap during post-glacial periods (i.e. Bollering allerod, Henrich stadial, and mid-holocene - present), two alternative scenarios emerge. First, species exhibiting secondary contact during periods of low glaciation, increases the chance for hybridization. A phenomenon observed among vertebrate and invertebrate systems in freshwater (Johnson et al., 2024; Moore et al., 2015; Nugent et al., 2024) and terrestrial (Carpenter et al., 2022; Dupuis & Sperling, 2015; Gompert et al., 2010; Rubidge et al., 2014) settings across the northern hemisphere. A second alternative interpretation is model misspecification due to the disjointed and sparse occurrence records of *S. hineana*, which may have produced non-analogue environments when projected into paleo-time slices (Elith et al., 2010; Fitzpatrick et al., 2009). Within our continuous predictions within the LGM, suitable habitat for *S. hineana* is only present within the Northeast coast of Connecticut, Massachusetts, and Maine, however our binary maps show no suitable habitat. It is possible that distributions for *S. hineana* were much larger during the Pleistocene and Pliocene, however the failure of our model to extrapolate to these time periods is a product of low sampling. Prospects of future research would be to estimate ancestral population sizes for the species to corroborate larger relictual distributions during interglacial periods.

### Genetic Analysis

Our phylogenetic analysis supports the validity of *S. hineana* and *S. tenebrosa* with high support, however we observe minimal phylogeographic structuring of either species both across nuclear and mitochondrial genes. Walker et al. (2020) shows, however, that a high degree of structuring exists among *S. hineana*, where haplotype diversity increases northward from the Ozarks to Great Lakes regions. Although our analysis and Walker (2020) use the same genetic marker (i.e. COI), we accredit this discrepancy due to the scope of our research including the range of our sampling and number of individuals. As for *S. tenebrosa*, it may have maintained gene flow across its range, similar to its circumpolar relative *Somatochlora sahlbergi* (Kohli et al., 2018). High gene flow across heterogeneous landscapes and continents have also been observed in both high dispersing taxa such as the migratory global skimmer *Pantala flavescens* (Troast et al., 2016; Ware et al., 2022), or regionally endemic montane species like *Synthemis eustalacta* (Goodman et al., 2026). We note, though, that the minimal evidence of genetic structuring in either species may be a result of low sampling.

Mitochondrial COI haplotype analyses exhibit definitive signals of hybridization within putative *S. tenebrosa* hybrids (SOM 11 & 12) and our newly discovered potential individual (SOM159), consistent with the findings of Hernandez and Cognato (2024). We note that the observed hybrids are the result of ♀ *S. hineana x ♂ S. tenebrosa*, as evident by our haplotype network exhibiting *S. tenebrosa* individuals with signatures of maternal *S. hineana* COI. ABBA-BABA tests further characterize these hybrids as introgressions with *S. tenebrosa*, also demonstrating the viability of hybrid individuals. Though, the absence of the reciprocal pairing or hybrids introgressed with *S. hineana* raises the likelihood that both hybridization and introgression are unidirectional in favor of *S. tenebrosa*. Bluetail damselfly (*Ischnura*) research also shows that asymmetry in post and prezygotic barriers may favor one cross over another, where egg fertility, failure of the male to inseminate, the female impeding insemination, and incompatible genetalia contribute to a significantly stronger reproductive barrier between ♀ *I. graellsii* mated to *♂ I. elegans* than the reciprocal pairing (Sánchez-Guillén et al., 2012). Other studies of hybridization in insects demonstrate that asymmetry emerges due to a number of factors, including mate choice, egg viability, or mechanical compatibility (Havill et al., 2017; Hochkirch & Lemke, 2011; Purcell et al., 2016). While Hernandez and Cognato (2024) and ourselves support the claim whereby differences in population size and asymmetrical mate pressure are the ultimate force of hybridization (Hubbs, 1955; Wirtz, 1999), we cannot discount the potential impact of additional prezygotic or postzygotic barriers that influence the direction of hybridization and introgression. Prospects for future research would involve assessing the compatibility of *♂ S. hineana x* ♀ *tenebrosa* or *S. hineana* x hybrids, as this would elucidate the strength *S. hineana*’s postzygotic barriers that are the result of inhabiting a specialized endemic habitat. Moreover, studying the ability of *S. hineana* individuals to compete for mates will test if contact with *S. tenebrosa* results in adaptive introgression (Arce-Valdés & Sánchez-Guillén, 2022) or genetic swamping (Todesco et al., 2016). As it currently stands, unidirectional introgression in favor of *S. tenebrosa* may threaten *S. hineana* populations with genetic swamping.

Our genetic and paleodistributional data suggest repeated instances of hybridization and introgression throughout the Pleistocene during interglacial periods. Post-glacial hybridization has also been recorded in both high and low-dispersing taxa including butterflies, chipmunks, fish, mussels, and birds (Carpenter et al., 2022; Dupuis & Sperling, 2015; Gompert et al., 2010; Johnson et al., 2024; Moore et al., 2015; Nugent et al., 2024; Rubidge et al., 2014). Within Odonata, most research pertaining to Quaternary hybridization and diversification focuses on the bluets (*Enallagma*). Reconstructed phylogenies of *Enallagma* point to Pleistocene glaciation sparking the rapid radiation of the genus, where numerous hybridization events are identified across clades (Turgeon et al., 2005). We hypothesize similar patterns of expansion and contraction are found in *S. hineana* and *S. tenebrosa,* where repeated secondary contact promoted increased incidences of hybridization among populations. Since the species diverged approximately ∼7ma (Goodman et al., 2025), along with *S. calverti* and *S. provocans*, divergence among this species complex was initially not driven by glaciation, yet hybridization potentially underwent during the glacial cycles of the Quaternary at ∼2.6mya (Goodman et al., 2025). Furthermore, prior hybridization events may have facilitated the maintenance of low prezygotic or postzygotic barriers and partial collapse of genetic distinctiveness (Xiong & Mallet, 2022) while still maintaining genetic species identities. Repeated contact and subsequent hybridization driven by glacial cycles have also been observed in other northern hemisphere glacial systems, including trees (Zhou et al., 2024). Future prospects for research will include sampling for hybrid specimens to examine potentially intermediate morphologies, as seen in other *Somatochlora* hybrids (Bick & Bick, 1981; S. Cannings & Cannings, 1985; Monroe & Britten, 2014; E. M. Walker, 1925), which would suggest the potential breakdown of prezygotic barriers.

## CONCLUSION

Our study finds evidence of hybridization occurring between *S. hineana* and *S. tenebrosa* among populations in Benzie County, Michigan as seen in Hernandez and Cognato (2024), while also identifying a new hybrid individual in Oregon County, Missouri. Minimal geographic structuring was recovered from our samples, though our analysis was limited in the number of individuals. Present-day distribution models of *S. hineana* and *S. tenebrosa* find areas of niche overlap that align with verified cooccurrences and paleo distribution models highlight potential hybridization events throughout the Pleistocene. By analysing hybrid individuals, we found evidence for unidirectional hybridization that may result in genetic swamping of the Hine’s Emerald. Therefore, we suggest that conservation efforts should focus on species complexes of varying degrees of hybridization as opposed to distinct taxonomic units.

## References

Abbott, J. C., Bota-Sierra, C. A., Guralnick, R., Kalkman, V., González-Soriano, E., Novelo-Gutiérrez, R., Bybee, S., Ware, J., & Belitz, M. W. (2022). Diversity of Nearctic dragonflies and damselflies (Odonata). Diversity, 14(7), 575.

Aiello-Lammens, M. E., Boria, R. A., Radosavljevic, A., Vilela, B., & Anderson, R. P. (2015). spThin: An R package for spatial thinning of species occurrence records for use in ecological niche models. Ecography, 38(5), 541–545.

Almendinger*, J. E., & Leete, J. H. (1998). Peat characteristics and groundwater geochemistry of calcareous fens in the Minnesota River Basin, USA. Biogeochemistry, 43(1), 17–41.

Almendinger, J. E., & Leete, J. H. (1998). Regional and local hydrogeology of calcareous fens in the Minnesota River Basin, USA. Wetlands, 18(2), 184–202.

Arce-Valdés, L. R., & Sánchez-Guillén, R. A. (2022). The evolutionary outcomes of climate-change-induced hybridization in insect populations. Current Opinion in Insect Science, 54, 100966.

Armstrong, M. R., Wilson, R. E., Johnson, J. A., Booms, T. L., Gesmundo, C. F., Pohlen, Z. M., Leonard, P. B., & Sonsthagen, S. A. (2025). Hybridization and Asymmetrical Introgression Between the Vulnerable Gray-Headed Chickadee and a More Abundant Congener, the Boreal Chickadee: Implications for Conservation. Ecology and Evolution, 15(7), e71673.

Bandelt, H.-J., Forster, P., & Röhl, A. (1999). Median-joining networks for inferring intraspecific phylogenies. Molecular Biology and Evolution, 16(1), 37–48.

Bick, G., & Bick, J. (1981). Heterospecific pairing among Odonata. Odonatologica, 10(4), 259–270.

Bonnald, J., Utge, J., Kuhner, M. K., Wasser, S. K., Asalu, E., Okimat, J. P., & Krief, S. (2021). Who are the elephants living in the hybridization zone? How genetics may guide conservation to better protect endangered elephants. Global Ecology and Conservation, 32, e01917.

Booth, T. H. (2022). Checking bioclimatic variables that combine temperature and precipitation data before their use in species distribution models. Austral Ecology, 47(7), 1506–1514.

Brown, J. L., Hill, D. J., Dolan, A. M., Carnaval, A. C., & Haywood, A. M. (2018). PaleoClim, high spatial resolution paleoclimate surfaces for global land areas. Scientific Data, 5(1), 180254.

Bybee, S., Córdoba-Aguilar, A., Duryea, M. C., Futahashi, R., Hansson, B., Lorenzo-Carballa, M. O., Schilder, R., Stoks, R., Suvorov, A., & Svensson, E. I. (2016). Odonata (dragonflies and damselflies) as a bridge between ecology and evolutionary genomics. Frontiers in Zoology, 13(1), 46.

Bybee, S. M., Kalkman, V. J., Erickson, R. J., Frandsen, P. B., Breinholt, J. W., Suvorov, A., Dijkstra, K.-D. B., Cordero-Rivera, A., Skevington, J. H., & Abbott, J. C. (2021). Phylogeny and classification of Odonata using targeted genomics. Molecular Phylogenetics and Evolution, 160, 107115.

Cannings, S., & Cannings, R. (1985). The larva of Somatochlora sahlbergi Trybŏm, with notes on the species in the Yukon Territory, Canada (Anisoptera: Corduliidae). Odonatologica, 14(4), 319–330.

Cannings, S. G., & Cannings, R. A. (1997). Dragonflies (Odonata) of the Yukon. Insects of the Yukon. Biological Survey of Canada (Terrestrial Arthropods), Ottawa, 169–200.

Capella-Gutiérrez, S., Silla-Martínez, J. M., & Gabaldón, T. (2009). trimAl: A tool for automated alignment trimming in large-scale phylogenetic analyses. Bioinformatics, 25(15), 1972–1973.

Carpenter, A., Graham, B., Spellman, G., & Burg, T. (2022). Do habitat and elevation promote hybridization during secondary contact between three genetically distinct groups of warbling vireo (Vireo gilvus)? Heredity, 128(5), 352–363.

Cashatt, E. D., & Vogt, T. E. (2001). Description of the larva of Somatochlora hineana with a key to the larvae of the North American species of Somatochlora (Odonata: Corduliidae). International Journal of Odonatology, 4(2), 93–105.

Catling, P. M. (2003). *Dragonflies (Odonata) of the Northwest Territories: Status ranking and preliminary atlas*. Northwest Territories, Environment and Natural Resources Ottowa, Canada.

Chunco, A. J. (2014). Hybridization in a warmer world. Ecology and Evolution, 4(10), 2019–2031.

Cognato, A. I., Seybold, S. J., & Sperling, F. A. (1999). Incomplete barriers to mitochondrial gene flow between pheromone races of the North American pine engraver, Ips pini (Say)(Coleoptera, Scolytidae). Proceedings of the Royal Society B: Biological Sciences, 266(1431), 1843.

Corbet, P. S. (1999). Dragonflies: Behaviour and ecology of Odonata.

Córdoba-Aguilar, A. (2008). Dragonflies and damselflies: Model organisms for ecological and evolutionary research. OUP Oxford.

Cozzolino, S., Nardella, A., Impagliazzo, S., Widmer, A., & Lexer, C. (2006). Hybridization and conservation of Mediterranean orchids: Should we protect the orchid hybrids or the orchid hybrid zones? Biological Conservation, 129(1), 14–23.

Craves, J. A., Cognato, A. I., O’Brien, D., & Mahoney, M. J. (2022). A new locality and unexpected haplotypes of the federally-endangered Hine’s Emerald dragonfly, Somatochlora hineana (Odonata: Corduliidae). Bulletin of American Odonatology, 13(2), 7.

Danecek, P., Bonfield, J. K., Liddle, J., Marshall, J., Ohan, V., Pollard, M. O., Whitwham, A., Keane, T., McCarthy, S. A., & Davies, R. M. (2021). Twelve years of SAMtools and BCFtools. Gigascience, 10(2), giab008.

Dijkstra, K.-D. B. (2025). Dragonflies and damselflies of the world: A guide to their diversity.

Doniol-Valcroze, P., Rancilhac, L., Brito, J., Miralles, A., Geniez, P., Benoit, L., Loiseau, A., Leblois, R., Dufresnes, C., & Crochet, P. (2024). Combining RAD_SEQ_ and contact zone analysis to decipher cryptic diversification in reptiles: Insights from the Spiny-footed Lizard (Reptilia, Lacertidae). Zoologica Scripta, 53(1), 1–15. 10.1111/zsc.12628

Dunkle, S. W. (2000). Dragonflies through binoculars: A field guide to dragonflies of North America. Oxford University Press.

Dupuis, J. R., & Sperling, F. A. (2015). Repeated reticulate evolution in North American Papilio machaon group swallowtail butterflies. PLoS One, 10(10), e0141882.

Elith, J., Kearney, M., & Phillips, S. (2010). The art of modelling range-shifting species. Methods in Ecology and Evolution, 1(4), 330–342.

Fitzpatrick, M. C., Preisser, E. L., Ellison, A. M., & Elkinton, J. S. (2009). Observer bias and the detection of low-density populations. Ecological Applications, 19(7), 1673–1679.

Foster, S., & Soluk, D. (2006). Protecting more than the wetland: The importance of biased sex ratios and habitat segregation for conservation of the Hine’s emerald dragonfly, Somatochlora hineana Williamson. Biological Conservation, 127(2), 158–166.

Gómez, J. M., González-Megías, A., Lorite, J., Abdelaziz, M., & Perfectti, F. (2015). The silent extinction: Climate change and the potential hybridization-mediated extinction of endemic high-mountain plants. Biodiversity and Conservation, 24(8), 1843–1857.

Gompert, Z., Lucas, L. K., Fordyce, J. A., Forister, M. L., & Nice, C. C. (2010). Secondary contact between Lycaeides idas and L. melissa in the Rocky Mountains: Extensive admixture and a patchy hybrid zone. Molecular Ecology, 19(15), 3171–3192.

Goodman, A., Abbott, J., Breinholt, J. W., Bybee, S., Frandsen, P. B., Guralnick, R., Kalkman, V. J., Kohli, M., Newton, L., & Ware, J. L. (2025). Systematics and biogeography of the Holarctic dragonfly genus Somatochlora (Anisoptera: Corduliidae). Systematic Entomology, 50(3), 585–610.

Goodman, A. M., Beatty, C. D., Büsse, S., Ubukata, H., Miyazaki, T., Blair, M. E., & Ware, J. L. (2024). Paleoecological niche modeling of Epiophlebia (Epiophlebioptera: Epiophlebiidae) reveals continuous distribution during the Last Glacial Maximum. International Journal of Odonatology, 27.

Goodman, A. M., Tolman, E. R., Beatty, C. D., Blair, M. E., Abbott, J. C., Alfaro, D. M., Bybee, S., Uche-Dike, R., Evangelista, D. A., & Frandsen, P. B. (2026). The Great Dividing Range as a driver of genetic divergence in a low-dispersing dragonfly. Ecological Entomology.

Goodman, A., Tolman, E., Uche-Dike, R., Abbott, J., Breinholt, J. W., Bybee, S., Frandsen, P. B., Gosnell, J. S., Guralnick, R., & Kalkman, V. J. (2023). Assessment of targeted enrichment locus capture across time and museums using odonate specimens. Insect Systematics and Diversity, 7(3), 5.

Grant, P. B., & Samways, M. J. (2007). Montane refugia for endemic and Red Listed dragonflies in the Cape Floristic Region biodiversity hotspot. Biodiversity and Conservation, 16(3), 787–806.

Guindon, S., Dufayard, J.-F., Lefort, V., Anisimova, M., Hordijk, W., & Gascuel, O. (2010). New algorithms and methods to estimate maximum-likelihood phylogenies: Assessing the performance of PhyML 3.0. Systematic Biology, 59(3), 307–321.

Harvey, J. A., Tougeron, K., Gols, R., Heinen, R., Abarca, M., Abram, P. K., Basset, Y., Berg, M., Boggs, C., & Brodeur, J. (2023). Scientists’ warning on climate change and insects. Ecological Monographs, 93(1), e1553.

Havill, N., Elkinton, J., Andersen, J., Hagen, S., Broadley, H. J., Boettner, G., & Caccone, A. (2017). Asymmetric hybridization between non-native winter moth, Operophtera brumata (Lepidoptera: Geometridae), and native Bruce spanworm, Operophtera bruceata, in the Northeastern United States, assessed with novel microsatellites and SNPs. Bulletin of Entomological Research, 107(2), 241–250.

Hayashi, F., Dobata, S., & Futahashi, R. (2005). Disturbed population genetics: Suspected introgressive hybridization between two Mnais damselfly species (Odonata). Zoological Science, 22(8), 869–881.

Hernandez, J., & Cognato, A. I. (2024). Incomplete barriers to heterospecific mating among Somatochlora species (Odonata: Corduliidae) as revealed in multi-gene phylogenies. Cladistics, 40(6), 598–617.

Hochkirch, A., & Lemke, I. (2011). Asymmetric mate choice, hybridization, and hybrid fitness in two sympatric grasshopper species. Behavioral Ecology and Sociobiology, 65(8), 1637–1645.

Hubbs, C. L. (1955). Hybridization between fish species in nature. Systematic Zoology, 4(1), 1–20.

Johnson, N. A., Henderson, A. R., Jones, J. W., Beaver, C. E., Ahlstedt, S. A., Dinkins, G. R., Eckert, N. L., Endries, M. J., Garner, J. T., & Harris, J. L. (2024). Glacial vicariance and secondary contact shape demographic histories in a freshwater mussel species complex. Journal of Heredity, 115(1), 72–85.

Kalyaanamoorthy, S., Minh, B. Q., Wong, T. K., Von Haeseler, A., & Jermiin, L. S. (2017). ModelFinder: Fast model selection for accurate phylogenetic estimates. Nature Methods, 14(6), 587–589.

Karger, D. N., Nobis, M. P., Normand, S., Graham, C. H., & Zimmermann, N. E. (2023). CHELSA-TraCE21k–high-resolution (1 km) downscaled transient temperature and precipitation data since the Last Glacial Maximum. Climate of the Past, 19(2), 439–456.

Kass, J. M., Muscarella, R., Galante, P. J., Bohl, C. L., Pinilla-Buitrago, G. E., Boria, R. A., Soley-Guardia, M., & Anderson, R. P. (2021). ENMeval 2.0: Redesigned for customizable and reproducible modeling of species’ niches and distributions. Methods in Ecology and Evolution, 12(9), 1602–1608.

Katoh, K., & Standley, D. M. (2013). MAFFT multiple sequence alignment software version 7: Improvements in performance and usability. Molecular Biology and Evolution, 30(4), 772–780.

Kawakami, T., & Butlin, R. K. (2012). Hybrid zones. Evolution & Diversity of Life, 1, 1–12.

Kessler, E. J., Mui, J. M., & Dreslik, M. J. (2026). Integrating multi-scale species distribution modeling and field validation to guide conservation of the endangered Hine’s Emerald Dragonfly (Somatochlora hineana). Journal of Insect Conservation, 30(3), 48.

Kohli, M. K., Sahlén, G., Kuhn, W. R., & Ware, J. L. (2018). Extremely low genetic diversity in a circumpolar dragonfly species, Somatochlora sahlbergi (Insecta: Odonata: Anisoptera). Scientific Reports, 8(1), 15114.

Kück, P., & Meusemann, K. (2010). FASconCAT: convenient handling of data matrices. Molecular Phylogenetics and Evolution, 56(3), 1115–1118.

Leigh, J. W., Bryant, D., & Nakagawa, S. (2015). POPART: full-feature software for haplotype network construction. Methods in Ecology & Evolution, 6(9), 1110.

Li, H., & Durbin, R. (2009). Fast and accurate short read alignment with Burrows–Wheeler transform. Bioinformatics, 25(14), 1754–1760.

Lobo, J. M., Jiménez-Valverde, A., & Real, R. (2008). AUC: a misleading measure of the performance of predictive distribution models. Global Ecology and Biogeography, 17(2), 145–151.

Malinsky, M., Matschiner, M., & Svardal, H. (2021). Dsuite-Fast D-statistics and related admixture evidence from VCF files. Molecular Ecology Resources, 21(2), 584–595.

Maresova, J., Habel, J. C., Neve, G., Sielezniew, M., Bartonova, A., Kostro-Ambroziak, A., & Fric, Z. F. (2019). Cross-continental phylogeography of two Holarctic Nymphalid butterflies, Boloria eunomia and Boloria selene. PLoS One, 14(3), e0214483.

Martin, S. H., Davey, J. W., & Jiggins, C. D. (2015). Evaluating the use of ABBA–BABA statistics to locate introgressed loci. Molecular Biology and Evolution, 32(1), 244–257.

Minh, B. Q., Schmidt, H. A., Chernomor, O., Schrempf, D., Woodhams, M. D., Von Haeseler, A., & Lanfear, R. (2020). IQ-TREE 2: New models and efficient methods for phylogenetic inference in the genomic era. Molecular Biology and Evolution, 37(5), 1530–1534.

Mirarab, S., & Warnow, T. (2015). ASTRAL-II: coalescent-based species tree estimation with many hundreds of taxa and thousands of genes. Bioinformatics, 31(12), i44–i52.

Monroe, E. M., & Britten, H. B. (2014). Conservation in Hine’s sight: The conservation genetics of the federally endangered Hine’s emerald dragonfly, Somatochlora hineana. Journal of Insect Conservation, 18(3), 353–363.

Moore, J., Bajno, R., Reist, J. D., & Taylor, E. B. (2015). Post-glacial recolonization of the North American Arctic by Arctic char (Salvelinus alpinus): Genetic evidence of multiple northern refugia and hybridization between glacial lineages. Journal of Biogeography, 42(11), 2089–2100.

Morales-Rozo, A., Tenorio, E. A., Carling, M. D., & Cadena, C. D. (2017). Origin and cross-century dynamics of an avian hybrid zone. BMC Evolutionary Biology, 17(1), 257.

Muhlfeld, C. C., Kovach, R. P., Jones, L. A., Al-Chokhachy, R., Boyer, M. C., Leary, R. F., Lowe, W. H., Luikart, G., & Allendorf, F. W. (2014). Invasive hybridization in a threatened species is accelerated by climate change. Nature Climate Change, 4(7), 620–624.

Muniz, A. C., Pimenta, R. J. G., Cruz, M. V., Rodrigues, J. G., Buzatti, R. S. de O., Heuertz, M., Lemos-Filho, J. P., & Lovato, M. B. (2022). Hybrid zone of a tree in a Cerrado/Atlantic Forest ecotone as a hotspot of genetic diversity and conservation. Ecology and Evolution, 12(1), e8540.

Naimi, B., & Naimi, M. B. (2017). Package ‘usdm.’ Uncertainty Analysis for Species Distribution Models. R-Cran, 18, 1–19.

Newton, L. G., Abbott, J. C., Bybee, S. M., Carter, P., Frandsen, P. B., Goodman, A., Guralnick, R., Hahn, B., Idec, J., & Kalkman, V. J. (2026). Soaring systematics: An evaluation of biogeography and flight behavior in dragonflies and damselflies (Insecta: Odonata) using phylogenomics. Systematic Biology, syag005.

Nugent, C. M., Kess, T., Langille, B. L., Beck, S. V., Duffy, S., Messmer, A., Smith, N., Lehnert, S. J., Wringe, B. F., & Kent, M. (2024). Post-glacial recolonization and multiple scales of secondary contact contribute to contemporary Atlantic salmon (Salmo salar) genomic variation in North America. Journal of Biogeography, 51(9), 1767–1782.

Owens, H., Barve, V., Chamberlain, S., Ram, K., Hart, T., & rOpenSci. (2025). *spocc: Interface to Species Occurrence Data Sources* (Version 1.2.4) [Computer software]. https://cran.r-project.org/web/packages/spocc/index.html

Phillips, S. J., Anderson, R. P., Dudík, M., Schapire, R. E., & Blair, M. E. (2017). Opening the black box: An open-source release of Maxent. Ecography, 40(7), 887–893.

Pintor, L. M., & Soluk, D. A. (2006). Evaluating the non-consumptive, positive effects of a predator in the persistence of an endangered species. Biological Conservation, 130(4), 584–591.

Purcell, J., Zahnd, S., Athanasiades, A., Türler, R., Chapuisat, M., & Brelsford, A. (2016). Ants exhibit asymmetric hybridization in a mosaic hybrid zone. Molecular Ecology, 25(19), 4866–4874.

Purdue, J., Gaige, T., Cashatt, E., & Vogt, T. (1999). Mitochondrial DNA sequence variation in populations of four species in the genus Somatochlora. Illinois State Museum, Springfield, Illinois. Pp.

R Core Team. (2021). A language and environment for statistical computing. R Foundation for Statistical Computing. 2017. Vienna, Austria: Available from: Http://Www.R-Project.Org/[Google Scholar].

Radosavljevic, A., & Anderson, R. P. (2014). Making better Maxent models of species distributions: Complexity, overfitting and evaluation. Journal of Biogeography, 41(4), 629–643.

Rubidge, E. M., Patton, J. L., & Moritz, C. (2014). Diversification of the Alpine Chipmunk, Tamias alpinus, an alpine endemic of the Sierra Nevada, California. BMC Evolutionary Biology, 14(1), 34.

Samways, M. J., Córdoba-Aguilar, A., Deacon, C., Alves-Martins, F., Baird, I. R., Barmentlo, S. H., Brasil, L. S., Bried, J. T., Clausnitzer, V., & Cordero-Rivera, A. (2025). Scientists’ warning on the need for greater inclusion of dragonflies in global conservation. Insect Conservation and Diversity, 18(4), 465–484.

Sánchez-Guillén, R. A., Córdoba-Aguilar, A., Hansson, B., Ott, J., & Wellenreuther, M. (2016). Evolutionary consequences of climate-induced range shifts in insects. Biological Reviews, 91(4), 1050–1064.

Sánchez-Guillén, R. A., Wellenreuther, M., & Cordero Rivera, A. (2012). Strong asymmetry in the relative strengths of prezygotic and postzygotic barriers between two damselfly sister species. Evolution: International Journal of Organic Evolution, 66(3), 690–707.

Sánchez-Guillén, Ra., Van Gossum, H., & Cordero Rivera, A. (2005). Hybridization and the inheritance of female colour polymorphism in two ischnurid damselflies (Odonata: Coenagrionidae). Biological Journal of the Linnean Society, 85(4), 471–481.

Schoener, T. W. (1968). The Anolis lizards of Bimini: Resource partitioning in a complex fauna. Ecology, 49(4), 704–726.

Shurtliff, Q. R. (2013). Mammalian hybrid zones: A review. Mammal Review, 43(1), 1–21.

Solano, E., Hardersen, S., Audisio, P., Amorosi, V., Senczuk, G., & Antonini, G. (2018). Asymmetric hybridization in Cordulegaster (Odonata: Cordulegastridae): Secondary postglacial contact and the possible role of mechanical constraints. Ecology and Evolution, 8(19), 9657–9671.

Taylor, S. A., Larson, E. L., & Harrison, R. G. (2015). Hybrid zones: Windows on climate change. Trends in Ecology & Evolution, 30(7), 398–406.

Todesco, M., Pascual, M. A., Owens, G. L., Ostevik, K. L., Moyers, B. T., Hübner, S., Heredia, S. M., Hahn, M. A., Caseys, C., & Bock, D. G. (2016). Hybridization and extinction. Evolutionary Applications, 9(7), 892–908.

Troast, D., Suhling, F., Jinguji, H., Sahlén, G., & Ware, J. (2016). A global population genetic study of Pantala flavescens. PloS One, 11(3), e0148949.

Turgeon, J., Stoks, R., Thum, R. A., Brown, J. M., & McPeek, M. A. (2005). Simultaneous Quaternary radiations of three damselfly clades across the Holarctic. The American Naturalist, 165(4), E78–E107.

Tynkkynen, K., Grapputo, A., Kotiaho, J. S., Rantala, M. J., Väänänen, S., & Suhonen, J. (2008). Hybridization in Calopteryx damselflies: The role of males. Animal Behaviour, 75(4), 1431–1439.

U.S. Fish and Wildlife Service. (2001). *Hine’s Emerald Dragonfly (Somatochlora hineana) Recovery Plan*. US Fish and Wildlife Service. https://ecos.fws.gov/docs/recovery_plan/hedplan.pdf

U.S. Fish and Wildlife Service. (2024). *Hine’s Emerald Dragonfly (Somatochlora hineana) 5-year Review*. US Fish and Wildlife Service. https://ecosphere-documents-production-public.s3.amazonaws.com/sams/public_docs/species_nonpublish/21792.pdf

Valavi, R., Guillera-Arroita, G., Lahoz-Monfort, J. J., & Elith, J. (2022). Predictive performance of presence-only species distribution models: A benchmark study with reproducible code. Ecological Monographs, 92(1), e01486.

Veloz, S. D. (2009). Spatially autocorrelated sampling falsely inflates measures of accuracy for presence-only niche models. Journal of Biogeography, 36(12), 2290–2299.

Vogt, T., & Cashatt, E. (1994). Distribution, habitat, and field biology of. *Somatochlora Hineana*.

Walker, E. M. (1925). The North American dragonflies of the genus Somatochlora (Issue 26). [Toronto] University of Toronto.

Walker, J., Mahoney, M., Templeton, A. R., McKenzie, P., Vogt, T. E., Cashatt, E. D., Smentowski, J., Day, R., Gillespie, R., & Henry, B. (2020). Contrasting Ozark and Great Lakes populations in the endangered Hines emerald dragonfly (Somatochlora hineana) using ecological, genetic, and phylogeographic analyses. Conservation Science and Practice, 2(3), e162.

Ware, K., Mendoza, T., Jinguji, H., & Sahlén, A. (2022). Suhling (2022): Evidence for widespread gene flow and migration in the Globe Skimmer dragonfly Pantala flaves-cens. Internafional Journal of Odonatology, 25, 43–55.

Warren, D. L., Glor, R. E., & Turelli, M. (2008). Environmental niche equivalency versus conservatism: Quantitative approaches to niche evolution. Evolution: International Journal of Organic Evolution, 62(11), 2868–2883.

Warren, D. L., Glor, R. E., & Turelli, M. (2010). ENMTools: A toolbox for comparative studies of environmental niche models. Ecography, 33(3), 607–611.

Warren, D. L., & Seifert, S. N. (2011). Ecological niche modeling in Maxent: The importance of model complexity and the performance of model selection criteria. Ecological Applications, 21(2), 335–342.

Wellenreuther, M., Muñoz, J., Chávez-Ríos, J. R., Hansson, B., Cordero-Rivera, A., & Sánchez-Guillén, R. A. (2018). Molecular and ecological signatures of an expanding hybrid zone. Ecology and Evolution, 8(10), 4793–4806.

White, E. L., Hunt, P. D., Schlesinger, M. D., Corser, J. D., & deMaynadier, P. G. (2014). A conservation status assessment of Odonata for the northeastern United States. New York Natural Heritage Program, Albany.

Wirtz, P. (1999). Mother species–father species: Unidirectional hybridization in animals with female choice. Animal Behaviour, 58(1), 1–12.

Xiong, T., & Mallet, J. (2022). On the impermanence of species: The collapse of genetic incompatibilities in hybridizing populations. Evolution, 76(11), 2498–2512.

Zhang, C., Rabiee, M., Sayyari, E., & Mirarab, S. (2018). ASTRAL-III: polynomial time species tree reconstruction from partially resolved gene trees. BMC Bioinformatics, 19(6), 153. 10.1186/s12859-018-2129-y

Zhou, Q., Karunarathne, P., Andersson-Li, L., Chen, C., Opgenoorth, L., Heer, K., Piotti, A., Vendramin, G. G., Nakvasina, E., & Lascoux, M. (2024). Recurrent hybridization and gene flow shaped Norway and Siberian spruce evolutionary history over multiple glacial cycles. Molecular Ecology, 33(17), e17495.

