## Supplementary figures and images for "Historical Areas of Hybridization in Two Striped Emerald Dragonfly Species"

### Supplemental Fig. S1

#### Maximum Likelihood

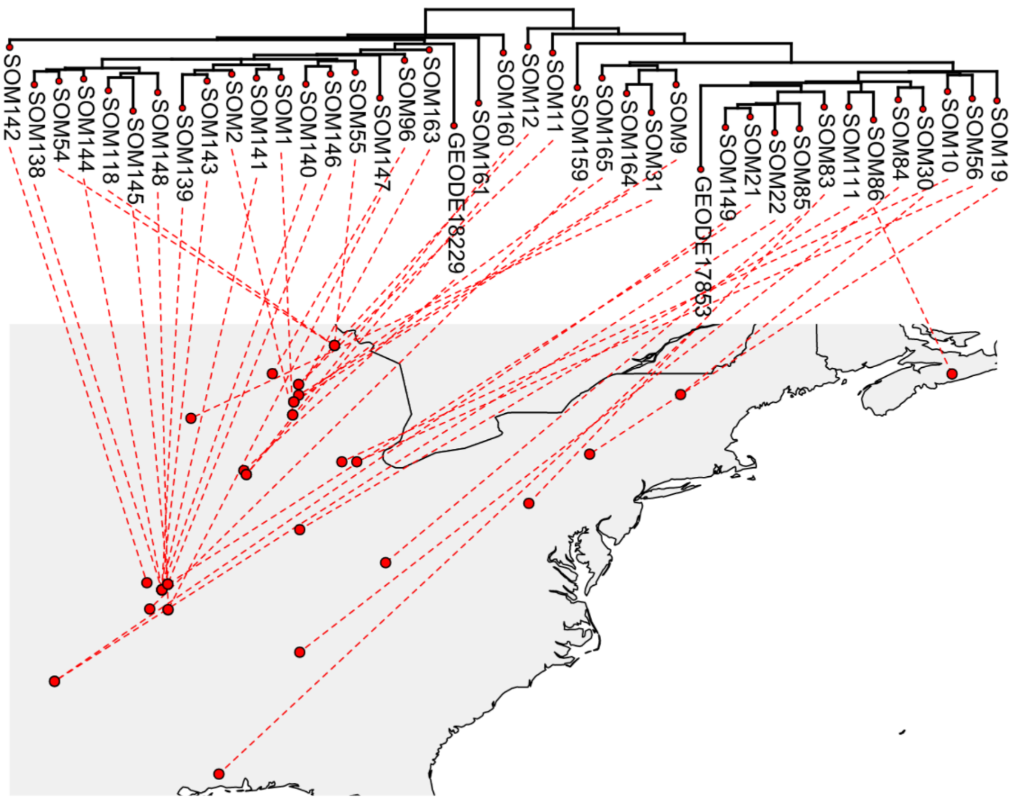


#### Astral

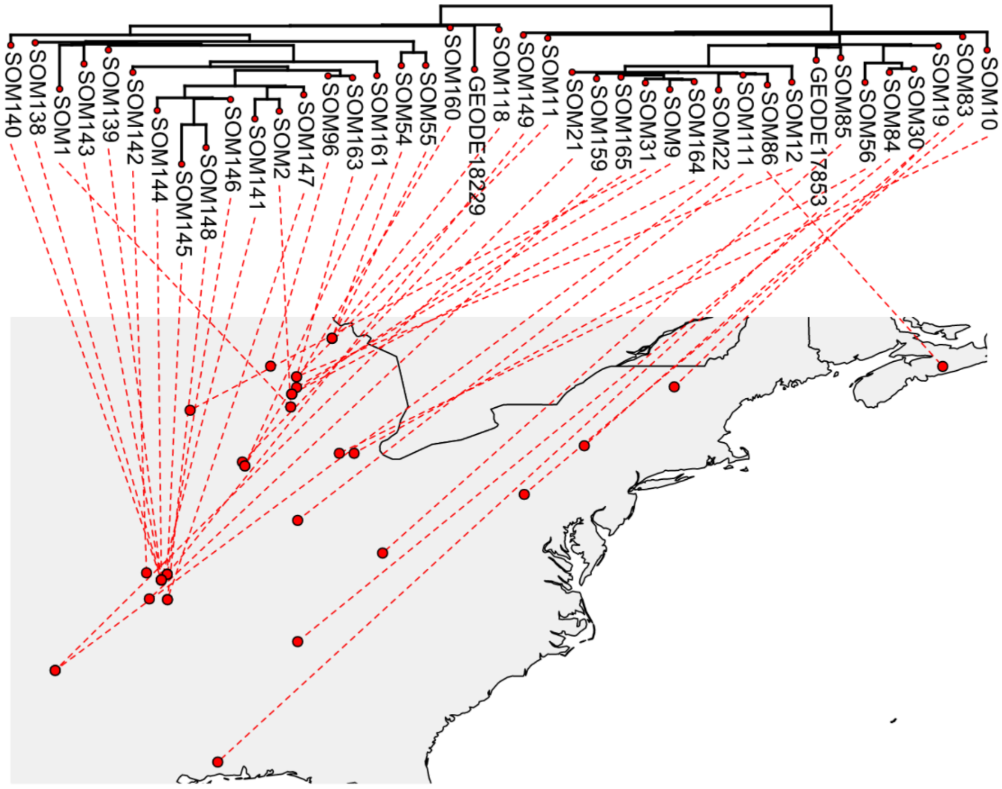

### Supplemental Fig. S2

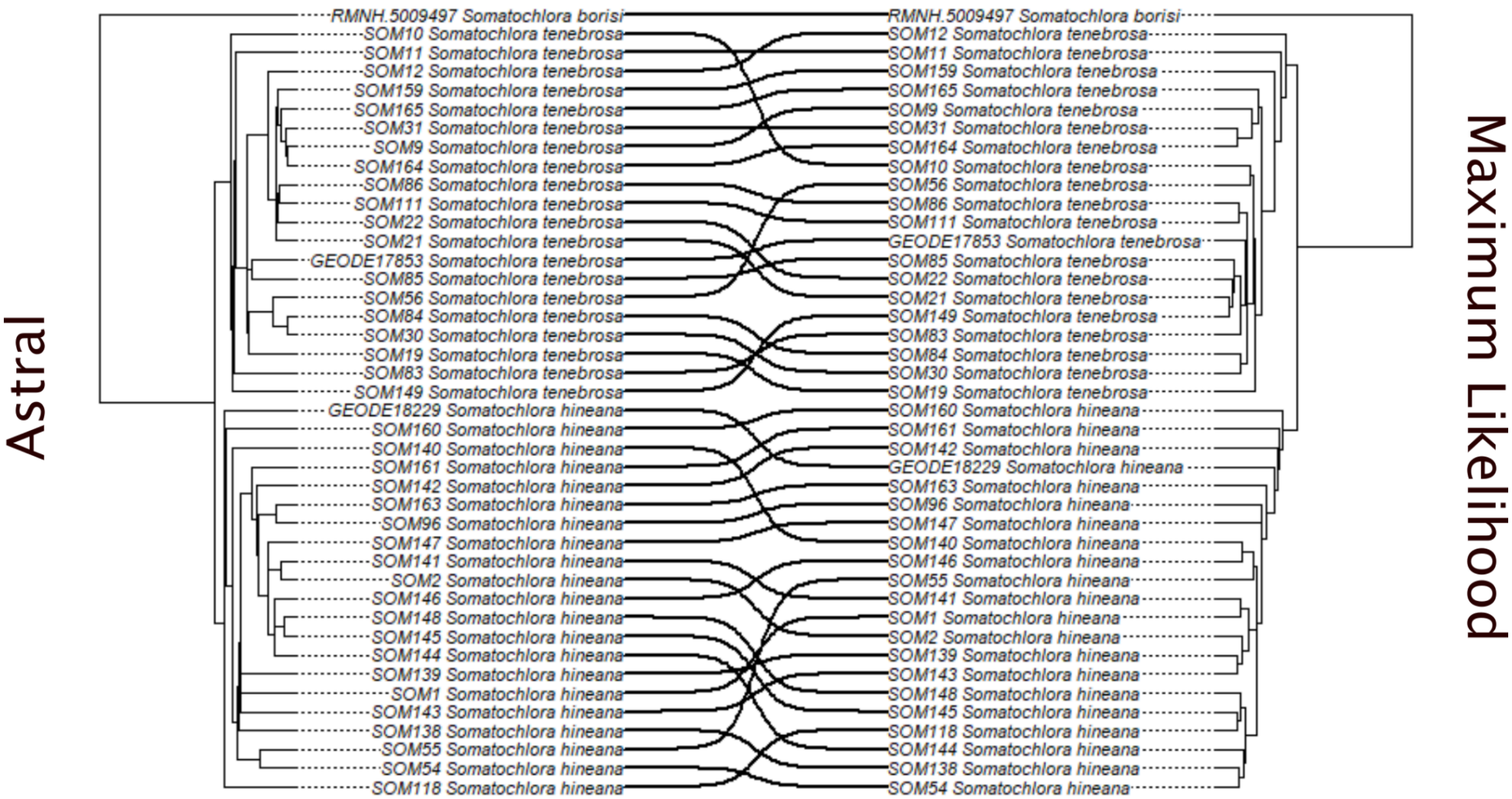

### Supplemental Fig. S3

*Somatochlora hineana*

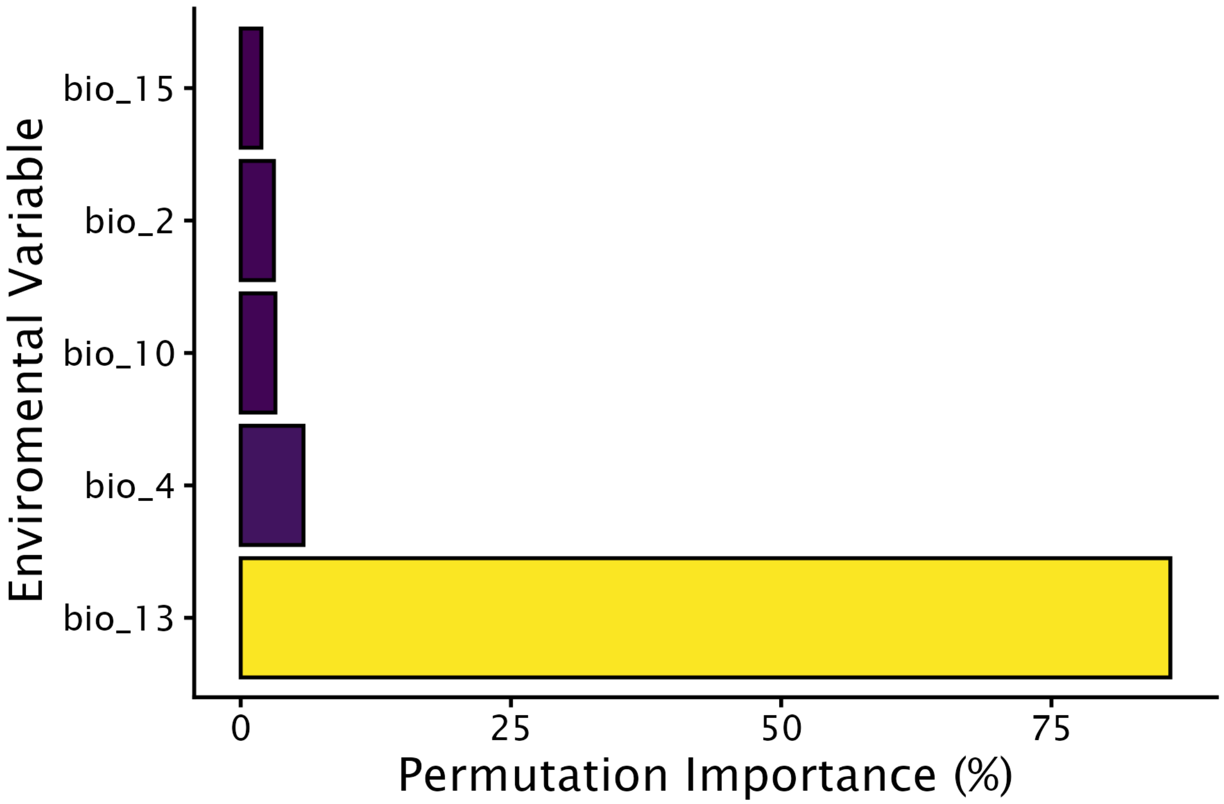


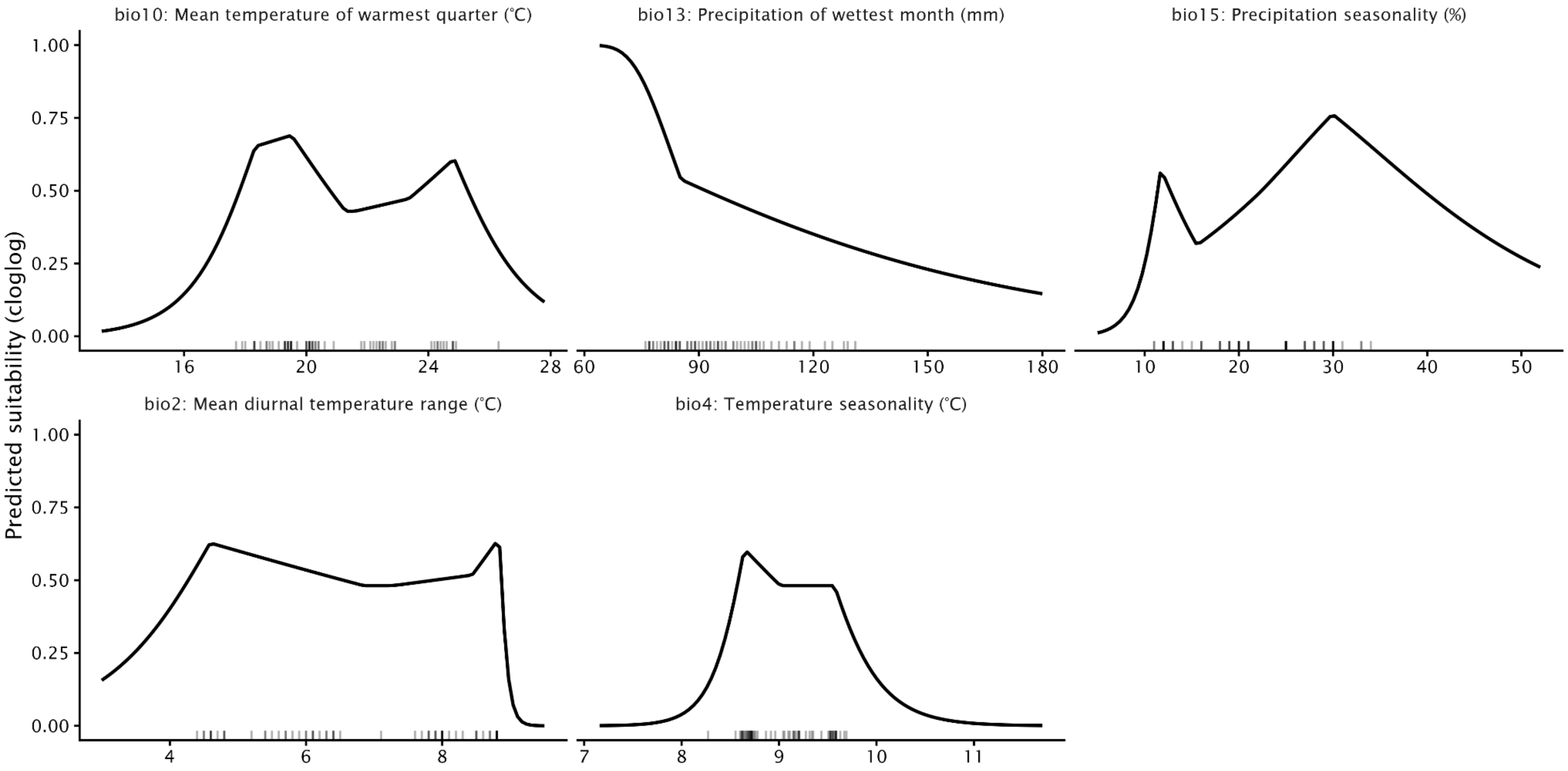

### Supplemental Fig. S4

*Somatochlora tenebrosa*

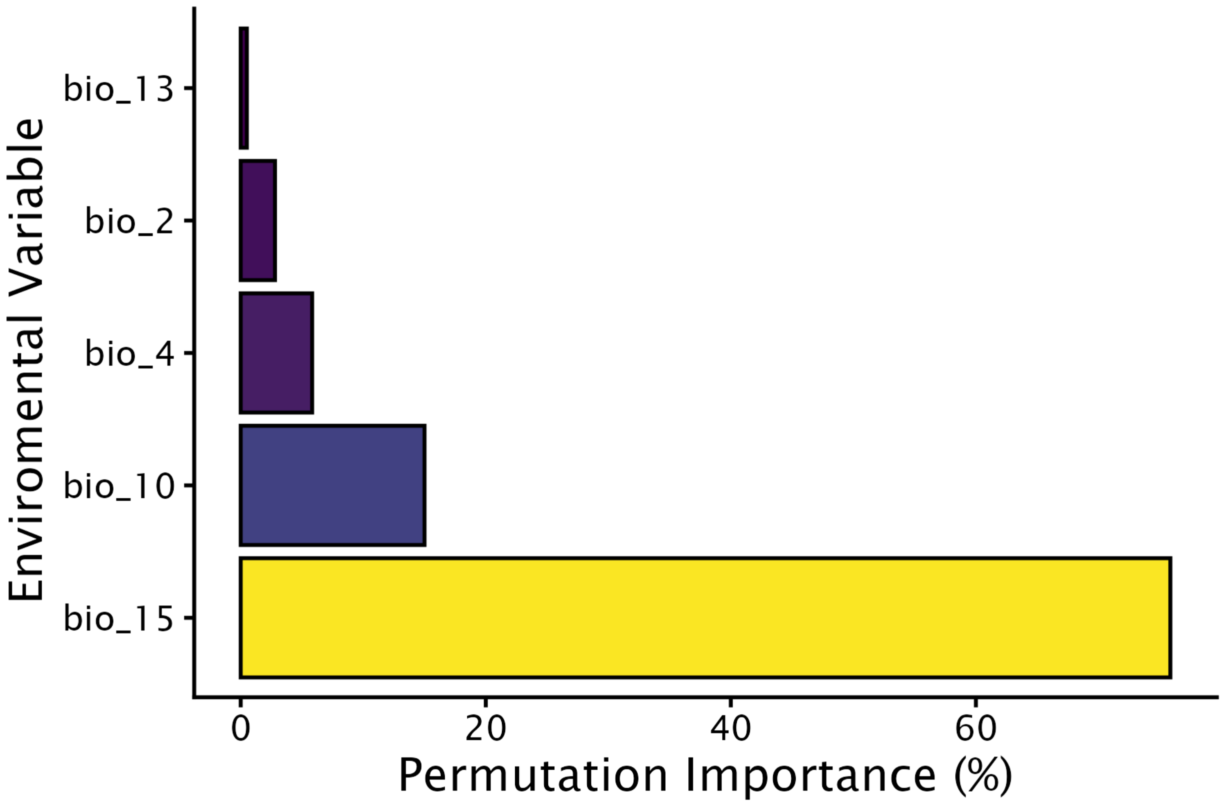


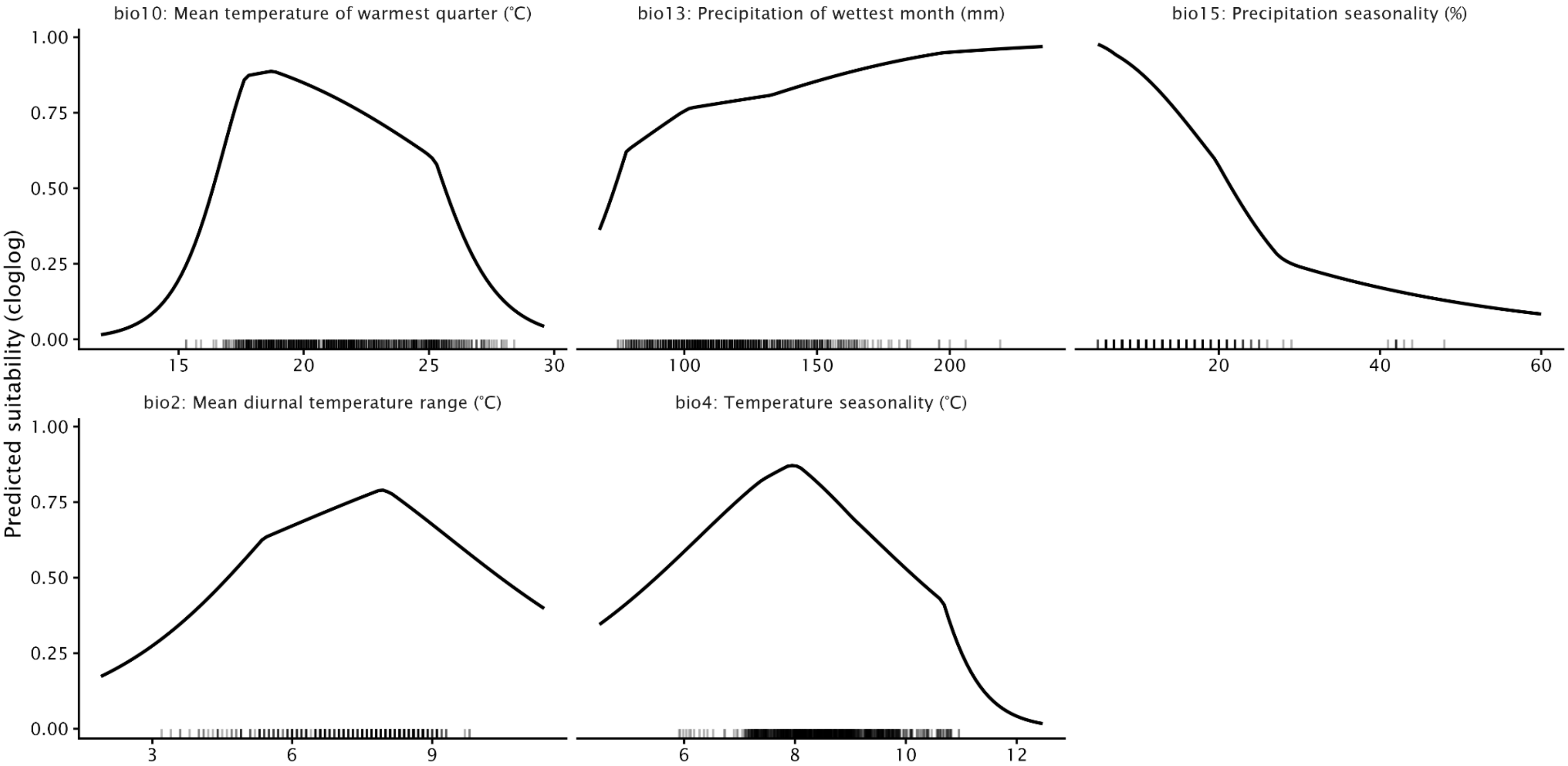
